# Identification of eight SARS-CoV-2 ORF7a deletion variants in 2,726 clinical specimens

**DOI:** 10.1101/2020.12.10.418855

**Authors:** Sun Hee Rosenthal, Ron M. Kagan, Anna Gerasimova, Ben Anderson, David Grover, Michael Hua, Yan Liu, Renius Owen, Felicitas Lacbawan

**Affiliations:** Quest Diagnostics Nichols Institute, San Juan Capistrano, CA 92675, USA; Quest Diagnostics Infectious Disease, San Juan Capistrano, CA 92675, USA

## Abstract

Severe acute respiratory syndrome coronavirus 2 (SARS-CoV-2) ORF7a, the ortholog of SARS-CoV ORF7a, is a type I transmembrane protein and plays an important role in virus-host interactions. Deletion variants in ORF7a may influence virulence, but only a few such isolates have been reported. Here, we report 8 unique ORF7a deletion variants of 6 to 96 nucleotides in length identified from 2,726 clinical specimens collected in March of 2020.

## Main Text

Although Severe acute respiratory syndrome coronavirus 2 (SARS-CoV-2) genomes are considered genetically stable, various mutations within the genome have been reported [1]. SARS-CoV-2 ORF7a, the ortholog of SARS-CoV ORF7a, is a type I transmembrane protein and plays an important role in virus-host interactions [2, 3]. Deletion variants in ORF7a may influence virulence, but only a few such isolates have been reported [4–6].

As part of an ongoing SPHERES Consortium project [7], we sequenced the SARS-CoV-2 genome from deidentified patient specimens submitted for testing at Quest Diagnostics in March 2020. Remnant extracted RNA of RT-PCR confirmed positive specimens was converted to cDNA and amplified with primers published by the ARTIC network [8]. Sequencing libraries were generated from the amplicons using the TWIST library preparation kit and sequenced on an illumina MiSeq (2×250 cycles). We utilized an in-house bioinformatics pipeline to generate consensus genomes and identified variants relative to the reference genome MN908947.3, Wuhan-Hu-1. Consensus genomes and sequencing reads are deposited under NCBI BioProject id PRJNA631061.

We identified 8 unique ORF7a deletion variants (Table 1) in 0.3% (8/2,726) clinical specimens. Specimens MW309829, MW309830, MW190280, and MW190429 had deletions in the ectodomain (96, 57, 18, and 12 nucleotides, respectively) associated with a loss of β-sheet strand(s) (Figure 1A) [9, 10]. In contrast to the previously reported ORF7a deletions [4–6], the deletions in these 4 isolates retained the signal peptide, transmembrane and cytoplasmic domains. The 96-nt and 57-nt deletion isolates also harbored ORF7a substitutions T120I or L102R. Specimens MW064508 and MW064714 harbored deletions in the transmembrane domain (9 and 6 nucleotides, respectively). Specimens MW309831 and MW314605 contained out-of-frame deletions of 40 and 13 nucleotides. The 40-nucleotide deletion resulted in the replacement of 18 amino acids spanning the transmembrane (partial) and cytoplasmic domains with 6 amino acids (KERQND, Figure 1A). The 13-nucleotide deletion removed the ORF7a stop codon and the ORF7b start codon, resulting in an extended ORF7a and a truncated ORF7b.

**Figure 1.**
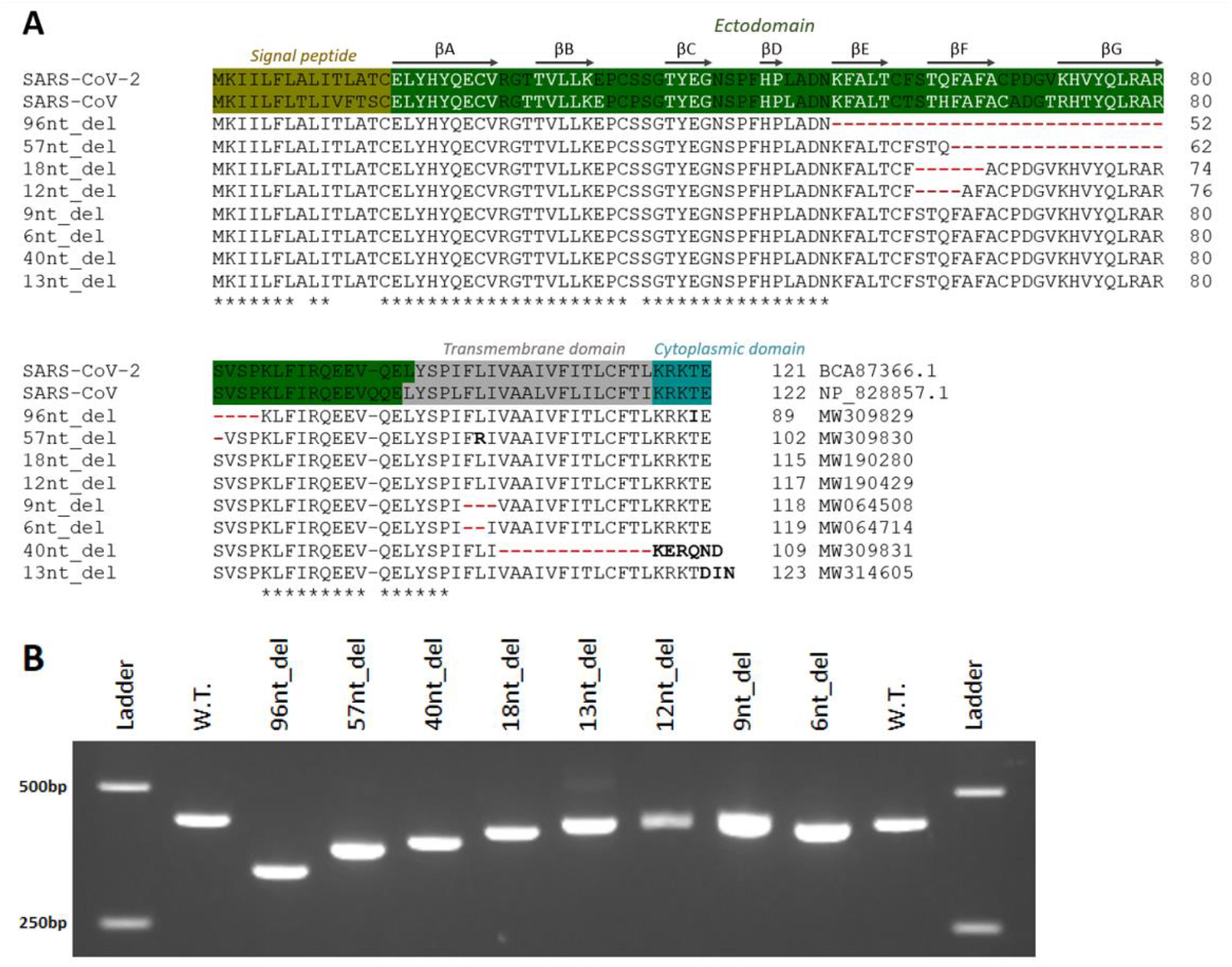
(A) ORF7a amino acid alignment of SARS-CoV-2, SARS-CoV, and 8 deletion isolates found in this study. The functional domains of ORF7a are indicated based on the previous reports [9, 10]. The arrows with labels denote the β-sheet strand regions reported by Zhou et al [10]. Deleted amino acids are represented by red lines; amino acid changes are marked in bold; and the conserved residues are shown as stars. (B) RT-PCR amplicons targeting the ORF7a-ORF7b regions. The expected amplicon size of the SARS-CoV-2 wild-type (W.T.) is 427bp, and the expected amplicon sizes of the deletion isolates of 96, 57, 40, 18, 13, 12, 9, and 6nt_del are 331, 370, 387, 409, 414, 415, 418, and 421bp, respectively.

**Table 1.** SARS-CoV-2 ORF7a deletion variants found in this study.

| NCBI Genbank accession/<br>SRA accession<sup>1</sup> | Lineage | Ct (N1)<sup>2</sup> | Position | Deletion<br>length (nt) | Alt_depth<sup>3</sup> | Alt_freq<br>(%)<sup>4</sup> | Other amino acid substitutions |
| --- | --- | --- | --- | --- | --- | --- | --- |
| MW309829/<br>SRR13167425 | B.1 | 22 | 27549 | 96 | 1702 | 97 | ORF1a:T265I, ORF1b:P314L, S:D614G,<br>ORF3a:Q57H, ORF7a:T120I |
| MW309830/<br>SRR13167194 | A.3 | 17 | 27578 | 57 | 1311 | 97 | ORF1a:D75E, ORF1a:P971L, ORF1b:F1757L,<br>ORF7a:L102R, ORF8:V62L, ORF8:L84S |
| MW309831/<br>SRR13167497 | B.1.36 | 20 | 27701 | 40 | 440 | 98 | ORF1a:T265I, ORF1b:P314L, S:D614G,<br>ORF3a:Q57H |
| MW190280/<br>SRR13106941 | B.1.2 | 21 | 27567 | 18 | 179 | 99 | ORF1a:T265I, ORF1b:P314L, ORF1b:G662S,<br>S:D614G, ORF3a:Q57H, ORF8:S24L |
| MW314605/<br>SRR13177577 | B.1 | 17 | 27755 | 13 | 181 | 21 | ORF1a:T265I, ORF1b:P314L, S:D614G,<br>ORF3a:Q57H |
| MW190429/<br>SRR13163447 | B.1.2 | 27 | 27567 | 12 | 158 | 38 | ORF1a:T265I, ORF1b:P314L, S:D614G,<br>ORF3a:Q57H, ORF8:S24L |
| MW064508/<br>SRR12980369 | B.1 | 13 | 27690 | 9 | 358 | 18 | ORF1a:T265I, ORF1b:P314L, S:D614G,<br>ORF3a:Q57H |
| MW064714/<br>SRR13004389 | B.1 | 12 | 27691 | 6 | 229 | 98 | ORF1a:T265I, ORF1b:P314L, S:D614G,<br>ORF3a:Q57H |

<sup>1</sup>SRA: sequence read archive; <sup>2</sup>cycle threshold value for the N1 real time PCR detector;
<sup>3</sup>coverage depth for the deletion; <sup>4</sup>percent of mapped reads with the deletion.

All deletion variants were confirmed by Sanger sequencing using amplicons targeting the ORF7a-ORF7b regions (**Figure 1B**). Three of the 8 isolates were heterogenous, with deletion variant frequency between 18% and 38% as determined by next-generation sequencing (NGS) (**Table 1**). Phylogenetic lineage analysis showed that the 57-nt deletion isolate belonged to pangolin lineage A.3, and the remaining 7 isolates were lineage B.1 (n=4), B.1.2 (n=2), or B.1.36 (n=1) [11]. The RT-PCR test cycle thresholds of the 8 deletion isolates were between 12 and 27, indicating efficient viral replication.

A recent study found that SARS-CoV-2 ORF7a is an immunomodulating factor for human CD14+ monocytes triggering an aberrant inflammatory response; the 4-stranded (βA, βC, βF, and βG) β-sheet in the ectodomain was suggested to be the major functional interface [10]. As the 4 ectodomain deletion variants in our study resulted in the loss of at least 1 β-sheet strand within the functional interface, we hypothesize that those isolates would have lost ORF7a biochemical function. Further studies are needed to understand the effects of these deletions on viral fitness or the clinical course of COVID-19.

## Acknowledgments

We deeply thank the staff of the ARTIC Network for developing SARS-CoV-2 primers. The authors thank Kayla Livingston, Hansook Kim, Kevin Qu, Allan Acab, Dawn Shalhout, Lamela Umaru, and Andy Ruden for technical assistance and data collection; Laurence Bernstein for data processing; the staff of Molecular Genetics for sample collection and Sanger confirmation; and Jeff Radcliff, Michael Wagner, Pranoot Tanpaiboon, and Hema Kapoor for critical review of the manuscript. This research was supported by Quest Diagnostics.

## References

[1] van Dorp L, Acman M, Richard D, Shaw LP, Ford CE, Ormond L, Owen CJ, Pang J, Tan CCS, Boshier FAT, Ortiz AT, Balloux F. Emergence of genomic diversity and recurrent mutations in SARS-CoV-2. Infect Genet Evol. 2020 Sep;83:104351

[2] Tan YX, Tan TH, Lee MJ, Tham PY, Gunalan V, Druce J, Birch C, Catton M, Fu NY, Yu VC, Tan YJ. Induction of apoptosis by the severe acute respiratory syndrome coronavirus 7a protein is dependent on its interaction with the Bcl-XL protein. J Virol. 2007 Jun;81(12):6346–55.

[3] Taylor JK, Coleman CM, Postel S, Sisk JM, Bernbaum JG, Venkataraman T, Sundberg EJ, Frieman MB. Severe Acute Respiratory Syndrome Coronavirus ORF7a Inhibits Bone Marrow Stromal Antigen 2 Virion Tethering through a Novel Mechanism of Glycosylation Interference. J Virol. 2015 Dec;89(23):11820–33.

[4] Holland LA, Kaelin EA, Maqsood R, Estifanos B, Wu LI, Varsani A, Halden RU, Hogue BG, Scotch M, Lim ES. An 81-Nucleotide Deletion in SARS-CoV-2 ORF7a Identified from Sentinel Surveillance in Arizona (January to March 2020). J Virol. 2020 Jul 1;94(14):e00711–20.

[5] Joonlasak K, Batty EM, Kochakarn T, Panthan B, Kümpornsin K, Jiaranai P, Wangwiwatsin A, Huang A, Kotanan N, Jaru-Ampornpan P, Manasatienkij W, Watthanachockchai T, Rakmanee K, Jones AR, Fernandez S, Sensorn I, Sungkanuparph S, Pasomsub E, Klungthong C, Chookajorn T, Chantratita W. Genomic surveillance of SARS-CoV-2 in Thailand reveals mixed imported populations, a local lineage expansion and a virus with truncated ORF7a. Virus Res. 2020 Nov 20:198233.

[6] Addetia A, Xie H, Roychoudhury P, Shrestha L, Loprieno M, Huang ML, Jerome KR, Greninger AL. Identification of multiple large deletions in ORF7a resulting in in-frame gene fusions in clinical SARS-CoV-2 isolates. J Clin Virol. 2020 Aug;129:104523.

[7] SARS-CoV-2 Sequencing for Public Health Emergency Response, Epidemiology and Surveillance (SPHERES), a new national genomics consortium to coordinate SARS-CoV-2 sequencing across the United States. https://covid-19bb.com/spheres-cdc

[8] https://github.com/artic-network/artic-ncov2019/tree/master/primer_schemes/nCoV-2019/V3

[9] Nelson CA, Pekosz A, Lee CA, Diamond MS, Fremont DH. Structure and intracellular targeting of the SARS-coronavirus Orf7a accessory protein. Structure. 2005 13:75–85.

[10] Zhou Z, Huang C, Zhou Z, Huang Z, Su L, Kang S, Chen X, Chen Q, He S, Rong X, Xiao F, Chen J, Chen S. 2020. Structural Insight Reveals SARS-CoV-2 Orf7a as an Immunomodulating Factor for Human CD14+ Monocytes. Available at SSRN: https://ssrn.com/abstract=3699795 or http://dx.doi.org/10.2139/ssrn.3699795

[11] Rambaut A, Holmes EC, O’Toole Á, Hill V, McCrone JT, Ruis C, du Plessis L, Pybus OG. A dynamic nomenclature proposal for SARS-CoV-2 lineages to assist genomic epidemiology. Nat Microbiol. 2020 Nov;5(11):1403–1407.

